# *Ruminococcus hollandia* sp. nov. and *Ruminococcus vasco* sp. nov., two novel starch-degrading *Ruminococcus* isolated from the rumen of Holstein dairy cattle

**DOI:** 10.64898/2026.07.10.737842

**Authors:** Kayla Calapa, Rachel Bock, Jordan Embree, Jolie LoBrutto, Mallory Embree

## Abstract

This study investigated the genomic and biochemical characteristics of two amylolytic microbial strains, NATIVEDY160^T^ (= JE7B6^T^ = NRRL B-68523^T^) and NATIVEDY161^T^ (= JL13D9^T^, = NRRL B-68524^T^) isolated from the rumen of healthy Holstein dairy cattle. Both strains are obligately anaerobic, non-motile, Gram positive, catalase-negative, and oxidase-negative. Morphologically, NATIVEDY160^T^ grows in long coccoid chains while NATIVEDY161^T^ grows in short chains or pairs. NATIVEDY160^T^ can catabolize amygdalin, esculin/ferric citrate, and starch, compared to NATIVEDY161^T^ which utilizes amygdalin, arbutin, esculin/ferric citrate, glycogen, and D-maltose as determined by API 50 CH carbon panels. Starch degradation ability was verified for both strains, but neither showed cellulolytic activity as confirmed by starch agar and Congo red agar assays, respectively. HPLC analysis revealed that lactate was the primary end product of both strains’ carbohydrate fermentation, while strain NATIVEDY161^T^ also produced small amounts of acetate. 16S rRNA sequences from both strains cluster with the *Oscillospiraceae* (formerly *Ruminococcaceae*) lineage *Ruminococcus* species, but average nucleotide identity of either strain compared to closely related *Ruminococcus* members was under the species threshold (95%). Genomic, phylogenetic, and phenotypic interrogation support NATIVEDY160^T^ and NATIVEDY161^T^ as novel species. Each strain was isolated from the rumen of dairy cows located within the central valley of southern California, which has a rich history of Dutch and Basque dairy farm ownership and is still the case today in the region. In recognition of the contributions and heritage of the central and southern California dairy industry, the names *Ruminococcus hollandia* and *Ruminococcus vasco* are proposed with NATIVEDY160^T^ and NATIVEDY161^T^ as their respective type strains.

---

The rumen of dairy cattle is an ecologically diverse and complex environment that is home to a variety of organisms such as fungi, protozoa, phage, archaea, and bacteria [1–2]. The prevalence of bacterial inhabitants has been estimated to be as high as 10^11^ cells/mL of rumen fluid, providing a variety of beneficial metabolites to their host organism [3–4]. Some ruminal microbes participate in the colonization and degradation of plant material present in the rumen due to their ability to break down cellulose, complex carbohydrates and resistant starches that the animal is incapable of degrading on its own [5–9]. The relationship between host and microbial community is a mutualistic one; during the process of fermenting released sugars, these microorganisms produce volatile fatty acids which, in turn, help the host ruminant to meet their energy needs [7, 10–11].

The genus *Ruminococcus* is found in a wide variety of animal and environmental hosts (e.g. ruminant animals, human gut, soils, Greenlandic ice sheets), and genomic and meta-analyses have shown that *Ruminococcus* tend to be most numerous in animals with a plant-based diet, including dairy cows [11–12]. *Ruminococcus* species are described as being obligately anaerobic coccoid bacteria that depend upon fermentable carbohydrates for growth [13–14]. They are typically non-motile, stain Gram positive, and morphologically appear in chains or pairs [5]. A number of *Ruminococcus* strains have been found to degrade resistant starches by producing large hydrolytic enzyme complexes [1, 6, 7, 12].

The word *Ruminococcus* is a combination of both “ruminalis” (meaning “of the rumen”) as well as the Greek word “kokkos,” which can be translated as “grain” or “berry” [15]. The genus name was proposed in 1948 when Sijpesteijn isolated the type strain *Ruminococcus flavefaciens* from cattle rumen content [16]. The *Ruminococcus* genus is polyphyletic, primarily incorporating the families *Oscillospiraceae* (formerly *Ruminococcaceae*) and *Lachnospiraceae* [5, 17]. The *Oscillospiraceae* lineage that contains the Group 1 *Ruminococcus* strains and retains the genus name includes *Ruminococcus albus, Ruminococcus flavefaciens, Ruminococcus callidus, Ruminococcus champanellensis, Ruminococcus bovis*, and *Ruminococcus bromii* [7, 11, 18–20]. Group 2 *Ruminococcus* genera reside within the *Lachnospiraceae* lineage and many of these members have been taxonomically reclassified under either the genus *Blautia* or *Mediterraneibacter*. The *Lachnospiraceae* lineage includes *Mediterraneibacter (Ruminococcus) gnavus, Blautia (Ruminococcus) obeum, Blautia (Ruminococcus) hansenii, Blautia hydrogenotrophica* (*Ruminococcus hydrogenotrophicus*), *Blautia (Ruminococcus) schinkii*, and *Blautia (Ruminococcus) luti* [19–21]. True *Ruminococcus* species are widely accepted as being members of the *Oscillospiraceae* family and are more likely to test positive for the starch, cellulose, and fiber-degrading phenotypes [7, 11, 17].

In this study, we discuss the isolation, characterization, and phylogeny of two novel amylolytic Group 1 *Ruminococcus* strains from dairy cow rumen, NATIVEDY160^T^ and NATIVEDY161^T^.

## Isolation and Phenotypic Characterization

NATIVEDY160^T^ and NATIVEDY161^T^ were isolated from the rumen of Holstein dairy cows housed at a commercial dairy farm in Corcoran, CA. These strains were initially grown for 72 hours in unmodified reinforced clostridial media (RCM) broth (BD, Franklin Lakes, NJ) in an anaerobic environment (5% H_2_, 20% CO_2_, 75% N_2_) at 37°C. When isolated on unmodified anaerobic RCM solid media, NATIVEDY160^T^ and NATIVEDY161^T^ phenotypically appear as small, white, slightly opaque circular colonies with even margins. Like many of the *Ruminococcus* type strains, NATIVEDY160^T^ and NATIVEDY161^T^ are not motile, non-spore forming, catalase and oxidase negative, and obligately anaerobic as evidenced by the lack of growth in the presence of oxygen.

Microscopic visualization was performed using an Olympus BX43 light microscope with an Excelis MPX-5C Pro camera attachment and CaptaVision v 2.3.1.0 imaging software. In preparation for imaging, cells were grown in a custom reinforced clostridial broth medium, which consisted of 10.0 g/L peptone (Research Products International, Mount Prospect, IL), 10.0 g/L beef extract (Sigma-Aldrich, St. Louis, MO), 3.0 g/L yeast extract (Sigma-Aldrich, St. Louis, MO), 5.0 g/L dextrose (Nutricost, North Vineyard, UT), 5.0 g/L NaCl (Spectrum, New Brunswick, NJ), 1.0 g/L soluble starch, 0.5 g cysteine-HCl (Spectrum, New Brunswick, NJ), and 3.0 g/L sodium acetate (Sigma-Aldrich, St. Louis, MO). Media was separated into Balch tubes in 20 mL aliquots, sparged with 80% N_2_ / 20% CO_2_ [v/v] for 20 minutes, capped with a butyl stopper and aluminum crimp seal to maintain anaerobic conditions, and autoclaved at 121°C/15 psi for 20 minutes. Tubes were cooled to room temperature prior to adjusting pH to 7.0 with 4M NaOH prior to inoculation. Cells were grown at an incubation temperature of 37°C for 30 hours until reaching late log phase. Each strain was then visualized at 800-2000x magnification. NATIVEDY160^T^ present as small cocci (0.6-0.7 μm x 1.3-1.5 μm) and will commonly form long chains, while NATIVEDY161^T^ are small cocci (0.5-0.7 μm x 1.0-1.1 μm) and often forms short chains or doubles (Supplementary Figure 1). Gram stains were carried out as per the protocol described by Beveridge [22]. Results confirmed both strains stain Gram positive (Supplementary Figure 1).

### Genomic Characterization

Genomic DNA was extracted from pure cultures of NATIVEDY160^T^ and NATIVEDY161^T^ using the New England BioLabs (NEB) Monarch High Molecular Weight DNA Extraction Kit #T3060L (NEB, Ipswich, MA). Whole genome short-read libraries were prepared using a KAPA HyperPlus kit (Roche, Basel, CH) with the manufacturer’s recommended protocol and sequenced (1x300bp) on an Illumina MiSeq. Long read sequencing libraries were prepared using the Oxford Nanopore Technologies (ONT) Ligation v14 Sequencing Kit #SQK-LSK114 (ONT, Lexington, MA) following the ONT kit instructions. Whole genome long-read libraries were sequenced on a MinION Mk1B device using a FLO-MIN114 flow cell. Reads were basecalled with Dorado v 0.4.0 [23] and sequencing adapters were excised with Porechop v 0.5.1 [24]. Long reads consisting of less than 3500 bp were removed and all remaining reads quality checked with Filtlong v 0.2.1 [25] and Nanoplot v 1.41.6 [26], accordingly.

Quality-filtered Oxford Nanopore long-reads and Illumina short-reads were hybrid assembled using Unicycler v 0.5.0 [27]. The NATIVEDY160^T^ and NATIVEDY161^T^ final genome assemblies each produced one circular contig averaging greater than 1000X depth for long reads and greater than 60X for short reads as confirmed by Bedtools v 2.26.0 [28]. NATIVEDY160^T^ is 2,427,278 bp in size and contains 50.04% mol G-C content, while NATIVEDY161^T^’s chromosome was 2,475,787 bp in length and consists of 46.59% mol G-C content. A comparison of these two characteristics reveals NATIVEDY160^T^ and NATIVEDY161^T^ have genomes that are smaller than many other *Ruminococcus* species yet have higher G-C content, highlighting distinctions from other members of the genus (Table 1).

**Table 1.** Table adapted from 18. Genomic and fermentative characteristics comparison of NATIVEDY160^T^ and NATIVEDY161^T^ with other members of the *Ruminococcus genus*. Starch and cellulose degradation capabilities of *R. bovis*, NATIVEDY160^T^, and NATIVEDY161^T^ confirmed via starch agar and Congo red agar assays as described in the Physiology and Chemotaxonomy section. Biochemical capabilities and genomic information for all other strains obtained from literature [5, 20, 37–40].

| Characteristic | NATIVEDY160<sup>T</sup> | NATIVEDY161<sup>T</sup> | *R. bovis* | *R. bromii* | *R. albus* | *R. flavefaciens* | *R. champanellensis* | *R. callidus* | *R. lactaris* | *R. torques* |
| --- | --- | --- | --- | --- | --- | --- | --- | --- | --- | --- |
| Genome GC Content (%) | 50.0 | 46.6 | 34.6 | 41.0* | 44.7* | 46.9* | 53.3* | 49.1* | 42.7* | 41.7* |
| Genome Size (Mbp) | 2.43 | 2.48 | 2.44 | 2.15* | 3.71* | 2.70* | 2.54* | 3.10* | 2.73* | 3.00* |
| Major Fermentation Products (s) | L | A,L | A | A | A,F | A,F,S | A,S | A,S | A,F | A,L |
| Able to Ferment: |  |  |  |  |  |  |  |  |  |  |
| Arabinose | - | - | - | - | - | - | nd | - | - | - |
| Cellobiose | - | - | - | - | + | + | + | + | - | - |
| Cellulose | - | - | + | - | + | + | + | - | - | - |
| Glucose | - | - | + | + | + | - | - | + | + | + |
| Lactose | - | - | - | - | + | + | - | + | + | + |
| Mannose | - | - | - | w/- | + | - | - | - | w/- | w/- |
| Maltose | - | + | + | + | - | - | - | + | sd | w |
| Mannitol | - | - | - | - | - | - | nd | - | + | - |
| Raffinose | - | - | - | - | - | - | - | + | - | - |
| Starch | + | + | + | + | - | - | - | + | - | - |

Symbols: \* Average based on all assemblies in the NCBI database; sd, strain dependent; w, weak reaction; nd, no data; A, acetate; F, formate; L, lactate; S, succinate.

NATIVEDY160^T^ and NATIVEDY161^T^ full length 16S rRNA gene sequences were extracted from genome assemblies and run through the nucleotide Basic Local Alignment Search Tool (BLAST) against other publicly available sequences. The best matches for both strains were to unnamed *Ruminococcus sp*. The closest named 16S rRNA matches to NATIVEDY160^T^ and NATIVEDY161^T^ were to *Ruminococcoides bili* at 96.06% and 94.75% identity and *Ruminococcoides intestinale* at 95.69% and 95.75% identity, respectively. These values do not meet or exceed the minimum species threshold cutoff of 98.7% similarity [29].

NATIVEDY160^T^ and NATIVEDY161^T^ complete genomes were then computationally analyzed against GenBank type strain *Ruminococcus* species genomes and genome assemblies of close neighbors based on 16S rRNA sequence BLAST alignments (*Ruminococcoides bili, Ruminococcoides intestinale, Clostridium leptum, Caproicibacter fermentans, Hydrogeniiclostridium mannosilyticum, Solibaculum mannosilyticum*, and *Pseudoruminococcus massiliensis*). Pairwise comparison of genomes were analyzed using both BLAST and MUMmer Average Nucleotide Identity (ANI). The MUMmer algorithm has been shown to provide a better comparison for more closely related species (i.e. greater than 90% identity) [30]. As evidenced by Tables 2 & 3, while the percent identity values are above 90% in some cases, the percent coverage is incredibly low, most of which are less than a few percent, due to the genomes being too divergent. ANI genome relatedness is better captured in this case utilizing the ANI BLAST algorithm that indicated the closest neighbor to NATIVEDY161^T^ and NATIVEDY160^T^ were one another at approximately 78% identity and 40% coverage. The next closest matches to known *Ruminococcus* species were at 75% or less identity and less than 30% coverage, all of which are well below the 95% ANI species threshold cutoff (Tables 4 & 5) [31].

**Table 2.** Average Nucleotide Identity (ANI) of NATIVEDY160^T^ compared to genomically-related microorganisms and *Ruminococcus* genome assemblies using MUMmer. All NCBI GenBank accession numbers have been included for reference.

| Microorganism | GenBank Assembly # | ANI (%) | Coverage (%) |
| --- | --- | --- | --- |
| *Solibaculum mannosilyticum* 12CBH8<sup>T</sup> | GCA_015140235 | 96.7 | 0.643 |
| *Ruminococcus gauvreaui* DSM 19829<sup>T</sup> | GCA_000425525 | 94.8 | 0.382 |
| *Ruminococcus bovis* JE7A12<sup>T</sup> | GCA_005601135 | 93.0 | 1.590 |
| *Caproicibacter fermentans* EA1<sup>T</sup> | GCA_009746625 | 89.4 | 0.136 |
| *Ruminococcoides bili* IPLA60002<sup>T</sup> | GCA_023369675 | 88.8 | 1.850 |
| *Pseudoruminococcus massiliensis* Marseille P3876<sup>T</sup> | GCA_900291955 | 88.1 | 0.440 |
| *Hydrogeniiclostridium mannosilyticum* ASD2818<sup>T</sup> | GCA_003268275 | 87.9 | 0.358 |
| *Clostridium leptum* DSM 753<sup>T</sup> | GCA_000154345 | 86.2 | 0.316 |
| NATIVEDY161<sup>T</sup> | N/A | 86.1 | 16.800 |
| *Ruminococcus intestinalis* NSJ71<sup>T</sup> | GCA_014288065 | 85.6 | 1.360 |
| *Ruminococcus albus* DSM 20455<sup>T</sup> | GCA_000179635 | 85.2 | 0.222 |
| *Ruminococcus bromii* ATCC 27255<sup>T</sup> | GCA_002834225 | 84.9 | 2.160 |
| *Ruminococcus callidus* ATCC 27760<sup>T</sup> | GCA_000468015 | 84.8 | 0.240 |
| *Ruminococcus champanellensis* 18P13<sup>T</sup> | GCA_000210095 | 84.6 | 0.360 |
| *Ruminococcoides intestinale* CLAAAH216<sup>T</sup> | GCA_040095935 | 84.4 | 1.990 |
| *Ruminococcus flavefaciens* ATCC 19208<sup>T</sup> | GCA_000518765 | 84.4 | 0.276 |

**Table 3.** Average Nucleotide Identity (ANI) of NATIVEDY161^T^ compared to genomically-related microorganisms and *Ruminococcus* genome assemblies using MUMmer. All NCBI GenBank accession numbers have been included for reference.

| Microorganism | GenBank Assembly # | ANI (%) | Coverage (%) |
| --- | --- | --- | --- |
| *Solibaculum mannosilyticum* 12CBH8<sup>T</sup> | GCA_015140235 | 91.1 | 0.446 |
| *Ruminococcus bovis* JE7A12<sup>T</sup> | GCA_005601135 | 90.7 | 4.660 |
| *Ruminococcus gauvreauii* DSM 19829<sup>T</sup> | GCA_000425525 | 90.0 | 0.158 |
| *Hydrogenticlostridium mannosilyticum* ASD2818<sup>T</sup> | GCA_003268275 | 89.9 | 0.222 |
| *Caproicibacter fermentans* EA1<sup>T</sup> | GCA_009746625 | 89.5 | 0.136 |
| *Clostridium leptum* DSM 753<sup>T</sup> | GCA_000154345 | 89.3 | 0.304 |
| *Ruminococcus champanellensis* 18P13<sup>T</sup> | GCA_000210095 | 86.8 | 0.220 |
| *Ruminococcus callidus* ATCC 27760<sup>T</sup> | GCA_000468015 | 86.7 | 0.155 |
| *Pseudoruminococcus massiliensis* Marseille P3876<sup>T</sup> | GCA_900291955 | 86.2 | 0.553 |
| NATIVEDY160<sup>T</sup> | N/A | 86.1 | 15.100 |
| *Ruminococcoides bili* IPLA60002<sup>T</sup> | GCA_023369675 | 84.7 | 2.850 |
| *Ruminococcus albus* DSM20455<sup>T</sup> | GCA_000179635 | 84.5 | 0.271 |
| *Ruminococcus bromii* ATCC 27255<sup>T</sup> | GCA_002834225 | 84.1 | 4.260 |
| *Ruminococcoides intestinale* CLAAH216<sup>T</sup> | GCA_040095935 | 84.0 | 3.96 |
| *Ruminococcus flavefaciens* ATCC 19208<sup>T</sup> | GCA_000518765 | 84.0 | 0.360 |
| *Ruminococcus intestinalis* NSJ71<sup>T</sup> | GCA_014288065 | 84.0 | 2.840 |

**Table 4.** Average Nucleotide Identity (ANI) of NATIVEDY160^T^ compared to genomically-related microorganisms and *Ruminococcus* genome assemblies using BLAST. All NCBI GenBank accession numbers have been included for reference.

| Microorganism | GenBank Assembly # | ANI (%) | Coverage (%) |
| --- | --- | --- | --- |
| NATIVEDY161<sup>T</sup> | N/A | 78.1 | 42.50 |
| *Ruminococcus bromii* ATCC 27255<sup>T</sup> | GCA_002834225 | 73.9 | 27.00 |
| *Ruminococcus gauvreauii* DSM 19829<sup>T</sup> | GCA_000425525 | 73.7 | 2.33 |
| *Rminococcoides intestinale* CLA-AA-H216 | GCA_040095935 | 73.5 | 25.00 |
| *Ruminococcoides bili* IPLA60002<sup>T</sup> | GCA_023369675 | 73.4 | 21.50 |
| *Ruminococcus intestinalis* NSJ71<sup>T</sup> | GCA_014288065 | 72.7 | 21.40 |
| *Solibaculum mannosilyticum* 12CBH8<sup>T</sup> | GCA_015140235 | 72.7 | 6.22 |
| *Ruminococcus albus* DSM 20455<sup>T</sup> | GCA_000179635 | 72.1 | 3.75 |
| *Ruminococcus bovis* JE7A12<sup>T</sup> | GCA_005601135 | 72.1 | 11.40 |
| *Ruminococcus torques* ATCC 27756<sup>T</sup> | GCA_000153925 | 72.0 | 2.70 |
| *Ruminococcus lactaris* ATCC 29176<sup>T</sup> | GCA_025152405 | 71.5 | 2.94 |
| *Clostridium leptum* DSM 753<sup>T</sup> | GCA_000154345 | 71.4 | 6.76 |
| *Ruminococcus flavefaciens* ATCC 19208<sup>T</sup> | GCA_000518765 | 71.0 | 4.49 |
| *Pseudoruminococcus massiliensis* Marseille P3876<sup>T</sup> | GCA_900291955 | 70.9 | 8.17 |
| *Hydrogeniclostridium mannosilyticum* ASD2818<sup>T</sup> | GCA_003268275 | 70.9 | 5.84 |
| *Ruminococcus champanellensis* 18P13<sup>T</sup> | GCA_000210095 | 70.5 | 5.77 |

**Table 5.** Average Nucleotide Identity (ANI) of NATIVEDY161^T^ compared to genomically-related microorganisms and *Ruminococcus* genome assemblies using BLAST. All NCBI GenBank accession numbers have been included for reference.

| Microorganism | GenBank Assembly # | ANI (%) | Coverage (%) |
| --- | --- | --- | --- |
| NATIVEDY160<sup>T</sup> |  | 77.5 | 38.60 |
| *Ruminococcus bovis* JE7A12<sup>T</sup> | GCA_005601135 | 75.4 | 16.10 |
| *Ruminococcus bromii* ATCC 27255<sup>T</sup> | GCA_002834225 | 74.8 | 29.00 |
| *Ruminococcoides intestinale* CLA-AA-H216<sup>T</sup> | GCA_040095935 | 74.3 | 27.60 |
| *Ruminococcoides bili* IPLA60002<sup>T</sup> | GCA_023369675 | 74.0 | 22.50 |
| *Ruminococcus intestinalis* NSJ71<sup>T</sup> | GCA_014288065 | 73.4 | 24.20 |
| *Ruminococcus albus* DSM20455<sup>T</sup> | GCA_000179635 | 71.9 | 3.54 |
| *Pseudoruminococcus massiliensis* Marseille P3876<sup>T</sup> | GCA_900291955 | 71.5 | 8.88 |
| *Ruminococcus torques* ATCC 27756<sup>T</sup> | GCA_000153925 | 71.4 | 3.23 |
| *Ruminococcus flavefaciens* ATCC 19208<sup>T</sup> | GCA_000518765 | 71.3 | 3.96 |
| *Ruminococcus gauvreauii* DSM 19829<sup>T</sup> | GCA_000425525 | 71.2 | 1.97 |
| *Solibaculum mannosilyticum* 12CBH8<sup>T</sup> | GCA_015140235 | 71.2 | 5.33 |
| *Clostridium leptum* DSM 753<sup>T</sup> | GCA_000154345 | 71.1 | 6.07 |
| *Ruminococcus lactaris* ATCC 29176<sup>T</sup> | GCA_025152405 | 70.7 | 2.73 |
| *Hydrogeniiclostridium mannosilyticum* ASD2818<sup>T</sup> | GCA_003268275 | 70.7 | 5.24 |
| *Ruminococcus callidus* ATCC 27760<sup>T</sup> | GCA_000468015 | 70.3 | 3.30 |

### Phylogeny

Full length 16S rRNA gene sequences were obtained from the Ribosomal Database Project (RDP) for all members of the order Clostridiales. Sequence alignment was performed using MUSCLE multiple sequence aligner and sequence ends trimmed in MEGA 11 software, leaving approximately 1400 bp [32–33]. NATIVEDY160^T^ and NATIVEDY161^T^ 16S rRNA sequences were analyzed on a neighbor-joining phylogeny with 500 bootstrap replicates constructed using MEGA 11 [33]. The tree was annotated and visualized with FigTree v 1.4.4 [34].

Out of the over 500 extracted Clostridiales sequences, 35 sequences were selected for a smaller tree, including NATIVEDY160^T^ and NATIVEDY161^T^, the *Ruminococcoides* genomes that are the closest identifiable matches to NATIVEDY160^T^ and NATIVEDY161^T^ as determined by 16S rRNA gene sequence BLAST alignment, all *Ruminococcus* type strains of the *Oscillospiraceae* (*Ruminococcaceae*) lineage, *Lachnospiraceae* Group 2 type strains classified as *Ruminococcus* and those that have since been re-classified under the genus *Blautia* or *Mediterraneibacter*, and representatives of neighboring clades as determined by the original RDP sequence phylogeny. The final 16S rRNA gene cladogram was rooted using *Halobacteroides elegans* Z-7287^T^ as the outgroup and bootstrap values displayed as decimals were amended to branches. Legacy nomenclature for re-classified *Ruminococcus sp*. can be seen listed in parenthesis and all NCBI accession numbers were placed in brackets next to branch taxon (Figure 1).

**Figure 1.**
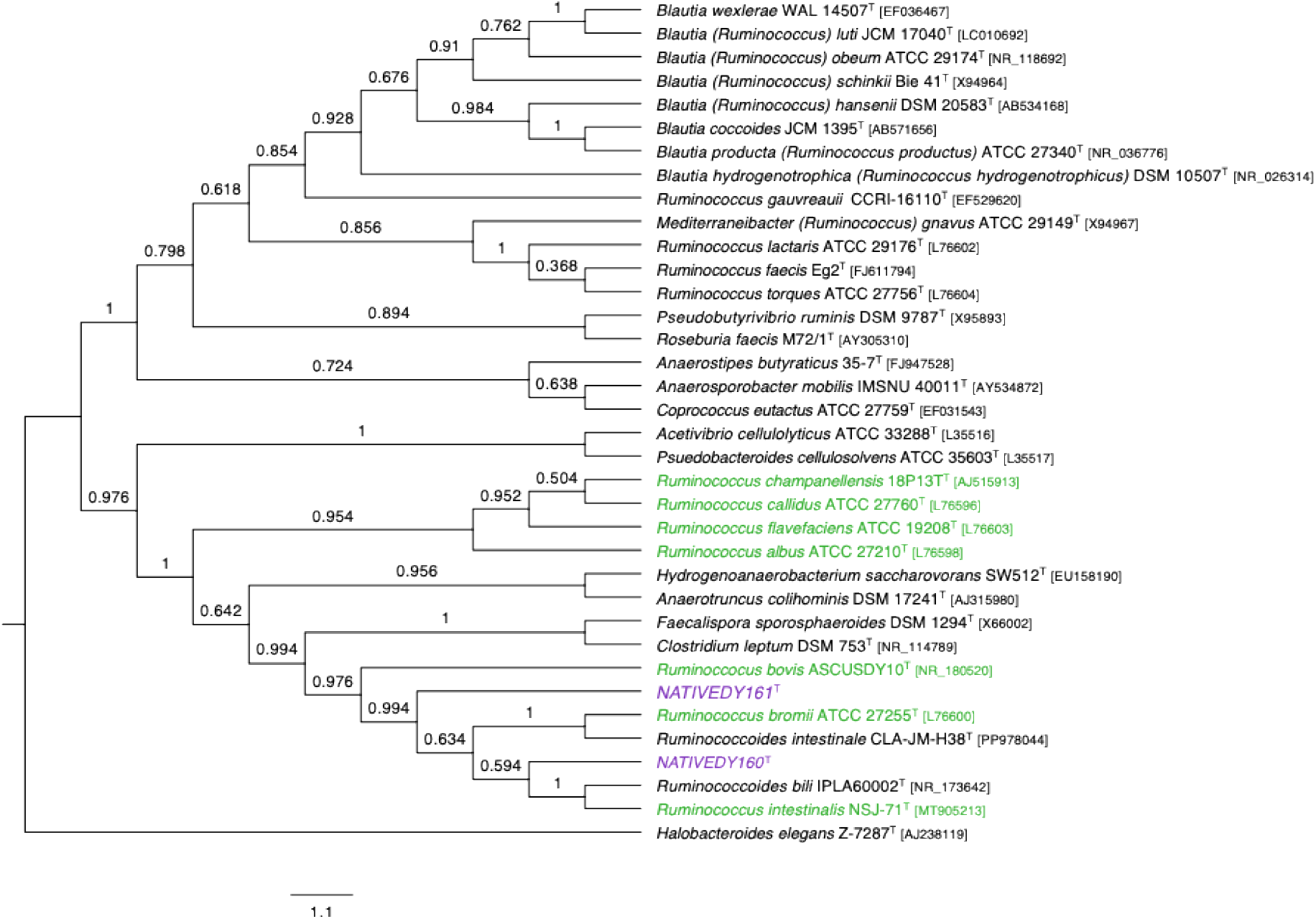
NATIVEDY160^T^ and NATIVEDY161^T^ 16S rRNA Gene Cladogram. 16S rRNA gene cladogram including NATIVEDY160^T^ and NATIVEDY161^T^, *Ruminococcus* type strains, members of the *Blautia* genus, *Mediterraneibacter* genus, and other close neighbors included in the *Lachnospiraceae* and *Oscillospiraceae* families. Cladogram was constructed in Mega 11 using MUSCLE multiple sequence aligner, and neighbor-joining phylogeny with 500 bootstrap replicates. The tree was visualized in FigTree, rooted using outgroup *H. elegans* Z-7287^T^, and bootstrap values added to each branch. NATIVEDY160^T^ and NATIVEDY161^T^ are labeled in purple, while members of the Group 1 *Ruminococcus sp*. are denoted in green. NCBI accession numbers were added in brackets next to taxon labels.

The Group 2 *Ruminococcus* (such as *R. torques* and *R. lactaris*) clustered with the *Mediterraneibacter sp*. and *Blautia sp*., while NATIVEDY160^T^ and NATIVEDY161^T^ were closer to the Group 1 *Ruminococcus* species such as *R. bromii, R. albus, R. bovis*, and *R. flavefaciens*. As expected, *R. intestinale, R. bromii*, and *R. bovis* were some of the species that clustered closest to NATIVEDY160^T^ and NATIVEDY161^T^ on the final tree, which appeared to align with the results of the ANI (Figure 1; Tables 4 & 5).

### Physiology and Chemotaxonomy

Catalase and oxidase confirmation was conducted using a 3% (v/v) hydrogen peroxide solution and 1.2% tetra-methyl-p-phenylenediamine solution, respectively, as outlined in the established protocols [35–36]. NATIVEDY160^T^ and NATIVEDY161^T^ exhibited negative results for both catalase and oxidase activity. To assess pH tolerance, strains were cultured in a custom anaerobic RCM broth medium (recipe as previously described). Twenty mL of broth was aliquoted into separate Balch tubes and pH adjusted in 0.5 increments from 4.0 to 9.0 with either NaOH or HCl post sparging and autoclaving as previously described. Cultures were incubated at 37°C for 48 hours and growth was estimated from optical density at 600 nm. Both strains demonstrated growth exclusively within the pH of 6.0-8.0, while the highest titers were observed at 7.0 and 7.5. No growth was observed at the pH ranges 4.0-5.5 and pH 8.5-9.0

Carbohydrate utilization profiles were determined using the API 50CH carbon panel (BioMérieux, Marcy-l’Étoile, France). Strains were first grown to late exponential phase in custom RCM broth, spun down at 16,000 x g for 3 mins, and supernatant removed. Cells were re-suspended in glucose and starch-free custom RCM broth medium and inoculated into honeycomb wells of the API test strips for 72 hours. One microliter of the pH indicator 0.425% phenol red was added to determine presence or absence of acid production.

NATIVEDY160^T^ was confirmed to catabolize amygdalin, esculin/ferric citrate, and starch and NATIVEDY161^T^ showed positive results in the presence of amygdalin, arbutin, esculin/ferric citrate, glycogen, and D-maltose (Supplementary Table 1).

Confirmation of starch utilization ability was performed using iodine agar stain assay as outlined in Lal, 2012 [37]. Media adapted for the assay consisted of custom RCM solid media with the starch substrate to test included as either soluble starch, starch from corn, or amylose from potato (Sigma-Aldrich, St. Louis, MO). NATIVEDY160^T^ and NATIVEDY161^T^ were streaked for isolation on all three plate types and incubated under anaerobic conditions at 37°C for 72 hours. Both strains were determined to be positive starch degraders, as verified by iodine staining after growth (Supplementary Figures 2 & 3).

Cellulose degradation assays were adapted from the protocol outlined by Szydlowski et al., 2015 [38]. To carry out the assay, starch was removed from the custom RCM solid media recipe, and media was subsequently amended with 0.2 g/L Congo red powder and 1.0 g/L either cellulose powder or carboxymethyl cellulose sodium salt (CMC) (Sigma-Aldrich, St. Louis, MO). Strains were streaked out and incubated under anaerobic conditions for 72 hours at 37°C prior to visualization with a Scan Interscience 500 (Interscience, Woburn, MA). Neither NATIVEDY160^T^ and NATIVEDY161^T^ tested positive for cellulolytic activity (Supplementary Figure 4).

Volatile fatty acid and metabolite production was verified by High Performance Liquid Chromatography (HPLC), using an Agilent 1260 Infinity II System with refractive index detection. Hi-Plex H column was set at 35°C at a flow rate of 0.2 mL/min, using a 5 mM sulfuric acid eluent. Both NATIVEDY160^T^ and NATIVEDY161^T^ produced lactate as their major fermentation end products and NATIVEDY161^T^ also produced a minor amount of acetate. Table 1 shows a comparison of fermentation products of NATIVEDY160^T^ and NATIVEDY161^T^ compared to other *Ruminococcus* species [5, 20, 37–40]. Notably, NATIVEDY160^T^ produces lactate alone, while other Group 1 strains produce acetate, succinate, and/or formate.

### Nomenclature and Description of *Ruminococcus hollandia* Sp. Nov. and *Ruminococcus vasco* Sp. Nov

The dairy industry in the United States, and particularly in the state of California, has ties to Basque, Portuguese, and Dutch history. Many Dutch and Basque dairymen settled in California’s San Joaquin Valley between the 1890s and 1950s [41–43]. During the 1950s, three areas (Dairyland, Cypress, and Dairy Valley) were the major dairy producing regions in California, but due to rising costs, dairymen in later decades expanded further into both Chino and San Joaquin Valleys [41, 44]. It is in these same areas of central and southern California that many dairies are still owned and operated by individuals of Dutch and Basque ancestry today: an area which is responsible for producing over 40 billion pounds of milk each year [41, 44].

*Ruminococcus* type strains NATIVEDY160^T^ and NATIVEDY161^T^ were isolated from the rumen of dairy cattle located in the San Joaquin Valley. We therefore propose the names *Ruminococcus hollandia* type strain NATIVEDY160^T^ and *Ruminococcus vasco* type strain NATIVEDY161^T^ to honor the significant contributions of the Basque and Dutch people on the dairy industry in California. The species name ‘*hollandia*’ pays homage to Dutch dairy farm history and ‘*vasco*’ pays tribute to the Basque dairy farm history.

Ruminococcus hollandia (hollandia L. m. gen. n. land of Holland)

*Ruminococcus hollandia* type strain NATIVEDY160^T^ is an obligately anaerobic, catalase/oxidase negative, Gram positive, non-motile, and non-spore-forming coccoid bacterium. Cells form long chains (0.6-0.7 μm x 1.3-1.5 μm) when grown in custom anaerobic RCM broth and colonies that form on custom RCM solid media appear small, white, slightly opaque, and circular. The closed genome for NATIVEDY160^T^ is 2,427,278 bp with 50.04% mol G-C content. NATIVEDY160^T^ is amylolytic, can utilize substrates such as amygdalin and esculin/ferric citrate as a carbon source, and produces lactate as the primary product of fermentation.

Type strain NATIVEDY160^T^ (= JE7B6^T^ = NRRL B-68523^T^) was isolated from rumen content of a healthy, Holstein cow from a dairy farm located in Corcoran, CA, USA.

*Ruminococcus vasco* (vasco L. m. gen. n. a Basque person)

*Ruminococcus vasco* type strain NATIVEDY161^T^ is an obligately anaerobic, catalase/oxidase negative bacterium. Cells (0.5-0.7 μm x 1.0-1.1 μm) stain Gram positive and form shorter coccoid chains or pairs when grown in a custom RCM broth. Colonies appear on custom RCM solid media as small, white, slightly opaque, and circular. The closed genome for NATIVEDY161^T^ is 2,475,787 bp in length and consists with 46.59% mol G-C content.

NATIVEDY161^T^ is amylolytic and can utilize amygdalin, D-maltose, glycogen, arbutin and esculin/ferric citrate as a carbon source. When cultured in custom RCM broth medium, the major fermentation products are lactate and acetate.

Type strain NATIVEDY161^T^ (= JL13D9^T^, = NRRL B-68524^T^) was isolated from rumen content of a healthy, Holstein cow from a diary farm located in Corcoran, CA, USA.

### Protologue

All sequences were deposited in the NCBI GenBank.

NATIVEDY160^T^ [JE7B6^T^] 16S rRNA Accession NCBI GenBank # PQ001355

NATIVEDY160^T^ [JE7B6^T^] WGS Accession NCBI GenBank # CP162391

NATIVEDY161^T^ [JL13D9^T^] 16S rRNA Accession NCBI GenBank # PQ001356

NATIVEDY161^T^ [JL13D9^T^] WGS Accession NCBI GenBank # CP162392

## AUTHOR STATEMENTS

### Ethical Statement

Sampling procedures were approved by veterinarians and the IACUC at Native Microbials, Inc.

## ABBREVIATIONS

ANI: Average nucleotide identity
RCM: Reinforced Clostridial Media
ONT: Oxford Nanopore Technologies
RDP: Ribosomal Database Project
CMC: carboyxymethyl cellulose sodium salt

