## Supplementary Materials for "*Ruminococcus hollandia* sp. nov. and *Ruminococcus vasco* sp. nov., two novel starch-degrading *Ruminococcus* isolated from the rumen of Holstein dairy cattle"

**Supplementary Figure 1. Methylene Blue and Gram Stains**

Methylene Blue stains for NATIVEDY160$^{T}$ (A) and NATIVEDY161$^{T}$ (B) and Gram stains for NATIVEDY160$^{T}$ (C) and NATIVEDY161$^{T}$ (D) after 72 hours of anaerobic incubation in custom RCM medium.

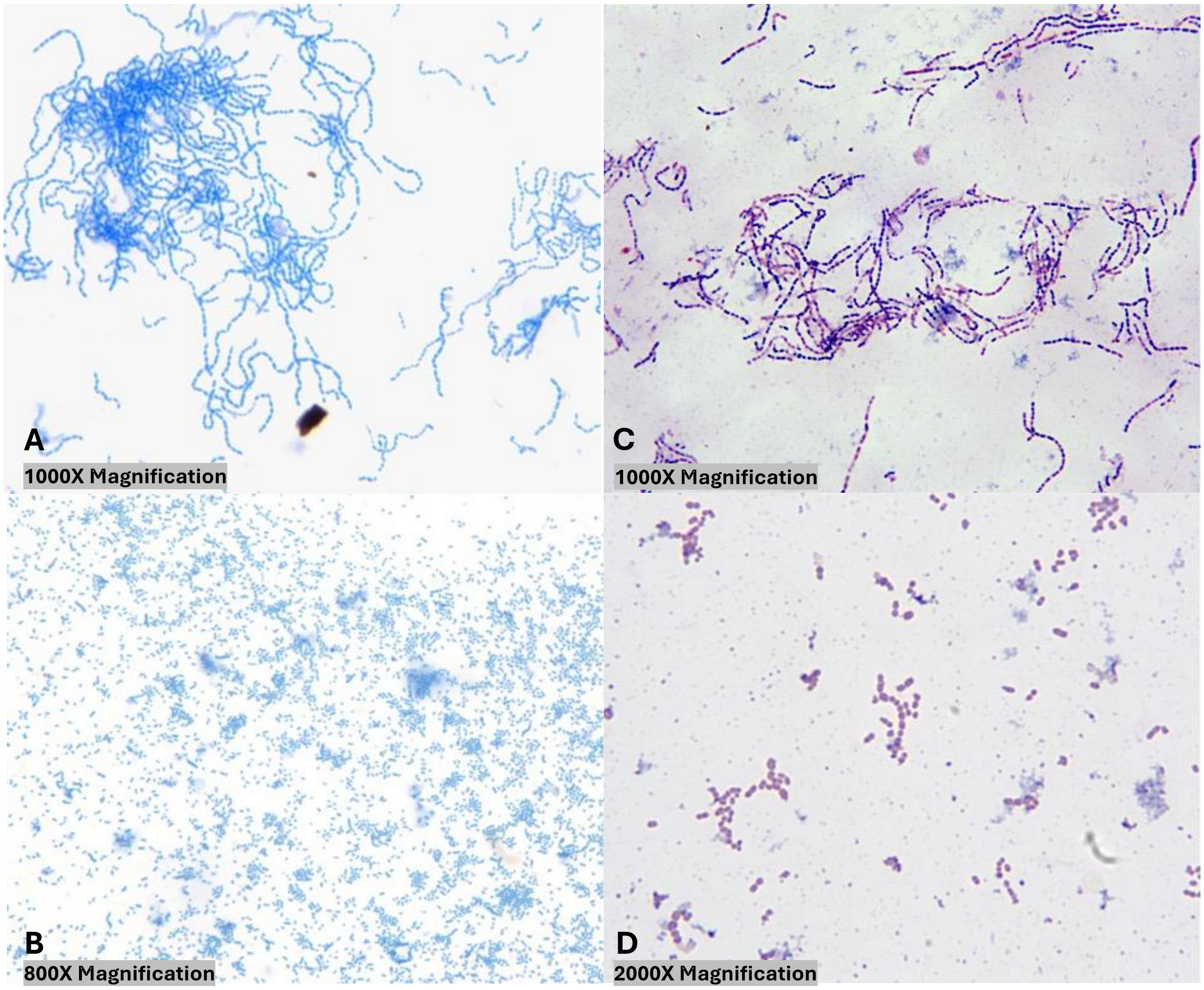

**Supplementary Table 1. API 50CH for NATIVEDY160$^{T}$ and NATIVEDY161$^{T}$**

Carbohydrate utilization profiles for NATIVEDY160$^{T}$ and NATIVEDY161$^{T}$ based on BioMérieux's API 50CH carbon panel.

| Component | NATIVEDY160$^{T}$ Growth (+/-) | NATIVEDY161$^{T}$ Growth (+/-) | Component | NATIVEDY160$^{T}$ Growth (+/-) | NATIVEDY161$^{T}$ Growth (+/-) |
|---|---|---|---|---|---|
| Control | - | - | Salicin | - | - |
| Glycerol | - | - | D-Cellobiose | - | - |
| Erythritol | - | - | D-Maltose | - | + |
| D-Arabinose | - | - | D-Lactose | - | - |
| L-Arabinose | - | - | D-Melibiose | - | - |
| D-Ribose | - | - | D-Saccharose | - | - |
| D-Xylose | - | - | D-Trehalose | - | - |
| L-Xylose | - | - | Inulin | - | - |
| D-Adonitol | - | - | D-Melezitose | - | - |
| Methyl-BD-xylopyranoside | - | - | D-Raffinose | - | - |
| D-Galactose | - | - | Starch | + | - |
| D-Glucose | - | - | Glycogen | - | + |
| D-Fructose | - | - | Xylitol | - | - |
| D-Mannose | - | - | Gentiobiose | - | - |
| L-Sorbose | - | - | D-Turanose | - | - |
| L-Rhamnose | - | - | D-Lyxose | - | - |
| Dulcitol | - | - | D-Tagatose | - | - |
| Inositol | - | - | D-Fucose | - | - |
| D-Mannitol | - | - | L-Fucose | - | - |
| D-Sorbitol | - | - | D-Arabitol | - | - |
| Methyl-aD-Mannopyranoside | - | - | L-Arabitol | - | - |
| Methyl-aD-Glucopyranoside | - | - | Potassium Gluconate | - | - |
| N-AcetylGlucosamine | - | - | Potassium 2-KetoGluconate | - | - |
| Amygdalin | + | + | Potassium 5-KetoGluconate | - | - |
| Arbutin | - | + | Esculin/Ferric Citrate | + | + |

**Supplementary Figure 2. NATIVEDY160$^T$ Streaked Out on Starch Media**

Bacterial starch utilization assay photos taken in natural lighting before (top row) and after (bottom row) iodine stain. NATIVEDY160$^T$ was grown in anaerobic conditions for 72 hours at 37°C. Various media types are indicated as follows: A = unstained soluble starch, B = iodine-stained soluble starch, C = unstained starch from corn, D = iodine-stained starch from corn, E = unstained amylose from potato, F = iodine-stained amylose from potato. A clearing within the purple iodine stain is indicative of a positive result.

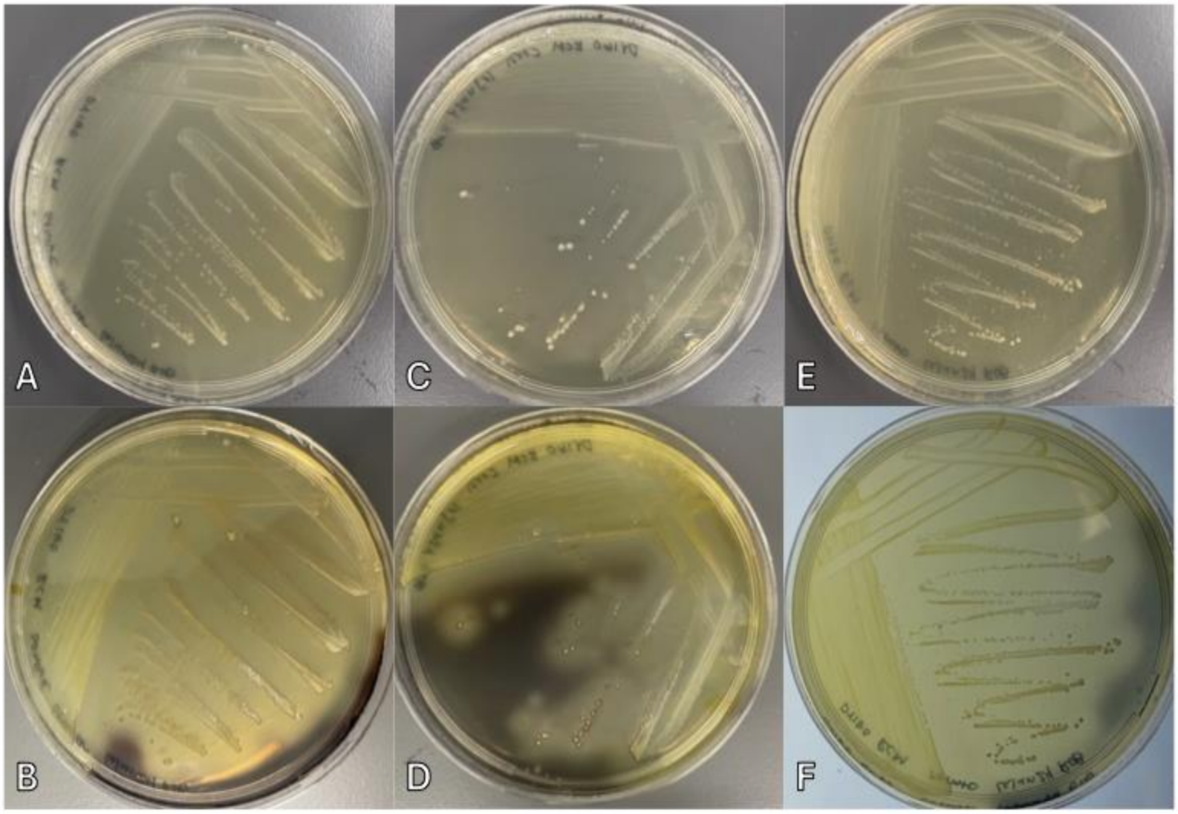

**Supplementary Figure 3. NATIVEDY161$^T$ Streaked Out on Starch Media**

Bacterial starch utilization assay photos taken in natural lighting before (top row) and after (bottom row) iodine stain. NATIVEDY161$^T$ was grown in anaerobic conditions for 72 hours at 37°C. Various media types are indicated as follows: A = unstained soluble starch, B = iodine-stained soluble starch, C = unstained starch from corn, D = iodine-stained starch from corn, E = unstained amylose from potato, F = iodine-stained amylose from potato. A clearing within the purple iodine stain is indicative of a positive result.

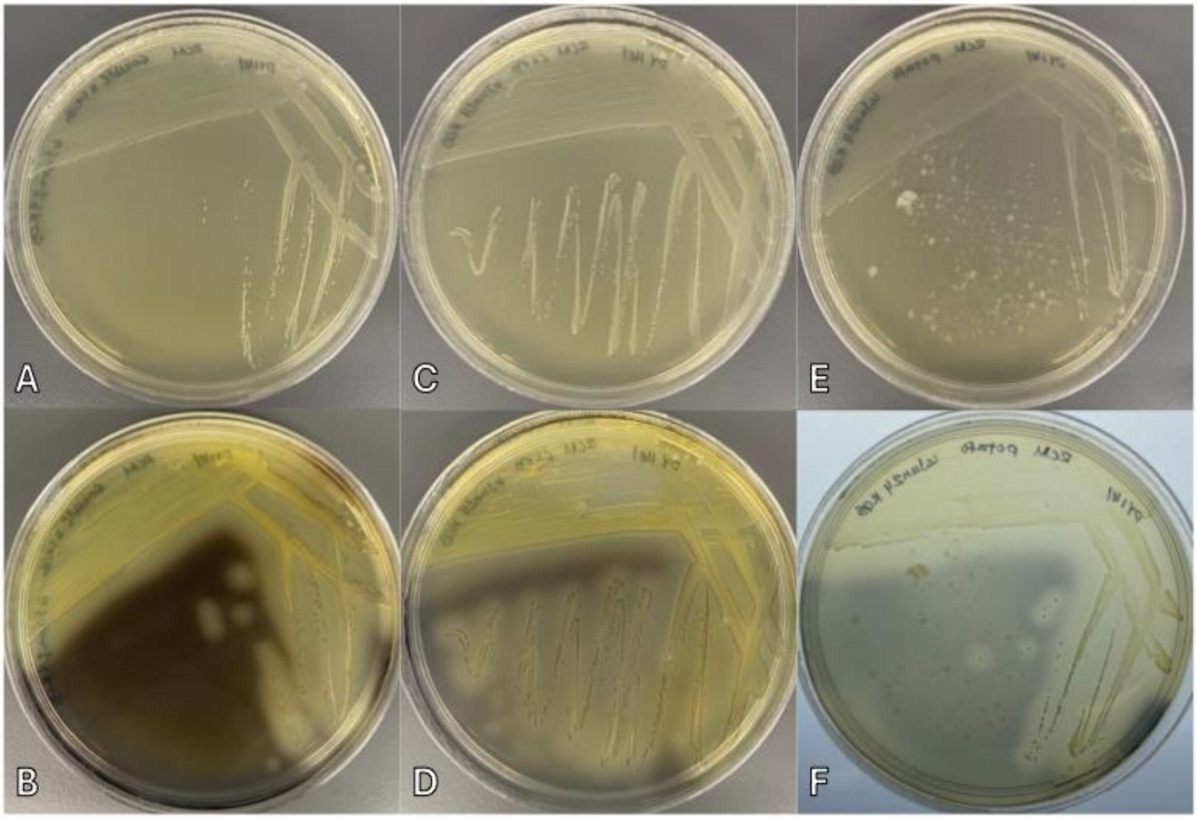

**Supplementary Figure 4. NATIVEDY160$^{T}$ and NATIVEDY161$^{T}$ Streaked Out on Cellulose or CMC Congo Red Media**

Cellulose degradation assay photos taken with backlighting using a Scan Interscience 500. NATIVEDY161$^{T}$ (bottom) and NATIVEDY160$^{T}$ (top) were grown in anaerobic conditions for 72 hours at 37°C on both custom RCM cellulose (left) and CMC (right) media. Clearing of the Congo red dye surrounding bacterial colonies would indicate positive result

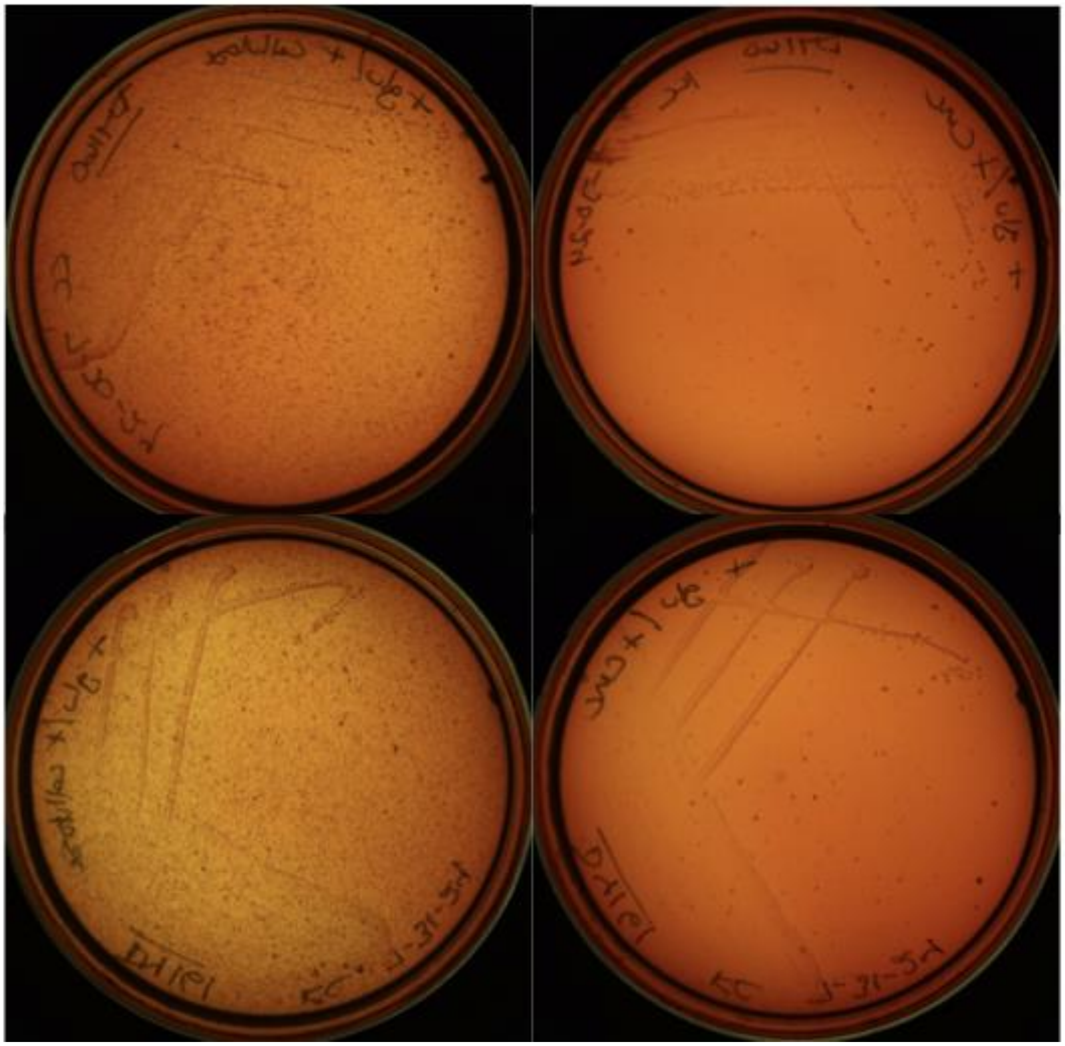
